# A Computational Model of Hippocampus: A Unified Theory About Engram and Sharp-Ware Ripples

**DOI:** 10.1101/2023.08.16.553536

**Authors:** ZHENG Zishuo

## Abstract

The hippocampus is key to memory encoding, consolidation, and retrieval. Previous work shows that neurons in the hippocampus fire in sequence to encode spatial information. The same group of cells will replay in memory consolidation, coupled with ripples, spindle, and slow waves. As for episodic memory, engram cells have been proposed to explain the encoding and transformation of episodic memory. Another universe theory about memory is the cognitive map theory. Here I use cognitive map theory as the bridge to overcome the gap between spatial and episodic memory. I believe spatial memory is a highly compressed case of episodic memory. In addition, I propose the hypothesis that engram can trigger sharp-wave ripples. I use a spike neural network-based computational model to verify this hypothesis. In conclusion, I believe engram cells and sharp-wave ripples are two different phenomena under a unified memory system.

## Background

### 1. Memory Medium - Engram Cells

When a new stimulus enters the brain, it is persisted and forms an underlying internal representation. Afterward, this representation can be extracted, consolidated, eliminated, and manipulated. Researchers try to find the physical medium for this process. Eventually, they discovered Engram. Memory engrams, also known as memory traces, were first proposed by Semon in his work. Semon defines memory engrams as “stimulated, persistent, latent modifications” [1]. Since then, the concept of engrams has been further enriched by other researchers. Tonegawa is a representative scholar in this field.

According to the previous work, engrams are considered the basic unit of memory. It is not equivalent to memory but is highly related to the formation of memory. More specifically, Engram cells refer to a cell population that satisfies the following conditions [2]: (1) Cells that fire frequently during the learning process. To identify such cells, biomarkers like c-Fos (an immediate early gene, IEG) can be used. (2) Cells that are changed by the learning process and undergoing chemical or physical changes, such as the excitability of cells, the density of dendrites and spines, and the strength of synapses. (3) By activating these cells with external stimuli or optogenetics, related memory recall can be triggered.

Engram cells have primarily been studied under the behavioral paradigm of episodic memory. A widely used behavioral paradigm for episodic memory is contextual fear conditioning (CFC) [3]. Engram cells are not only located in a specific brain region but are widely distributed across the brain. This property suggests that memories are not represented by a specific brain region but are distributed encoded. As for the hippocampus, engram cells that are related to episodic memory are found in the DG area. Existing experimental evidence about engrams can be divided into three levels, including observation [4–5], inhibition [6–9], and manipulation [10–13]. This evidence clarifies that engrams are indeed mediators of memory formation and representation.

### 2. Memory Encoding – Sequential Firing and Replay

Another story about memory representation is sequential firing. Previous findings proved that spatial memory is encoded by the sequential firing of neurons in the CA area. These cells are called place cells. Place cells always come up together with grid cells. They work like the reference frame and coordinates. When a mouse runs forward in a maze, the EC cortex will form a map of the maze with grid cells. And the mouse can locate its location with place cells in the hippocampus. As place cells are binding to a specific spatial location, they always fire in sequence to encode the time-varying coordinates of the mouse. That is how sequential firing comes into being.

During offline states such as sleep, information in the hippocampus is replayed. Place cells in the hippocampus will fire in the same pattern as the exploration and learning period. This is called replay. Replay is essential for memory consolidation which transfers information from the hippocampus into the isocortex. Experimental evidence shows that interrupting the replay process will damage learning performance.

To answer the question of how neurons in the hippocampus encode external information, Daoyun Ji devised a spatial memory task to answer this question [14]. Mice would run through a figure-of-eight maze, and the researchers recorded the activity of neurons in the CA1 region of the hippocampus. The results showed that neurons encode the spatial information of the maze by firing sequentially. Furthermore, during sleep, the same set of neurons fires in the same pattern, a phenomenon known as “sequential replay.” The researchers found that neuronal replay is highly coupled to sharp ripples, and replay often occurs within sharp ripples.

### 3. Memory Syntax - Sharp-wave Ripples

If engrams represent the basic unit of memory, the brain also needs grammar to organize these “memory letters”. This kind of grammar is called neural syntax, and its specific manifestation is brain rhythms or neural oscillation [16]. There are many different rhythmic oscillations in the brain, ranging from sharp-wave ripples, spindles, to slow waves. The mutual coupling between nerve rhythms of different frequencies creates a hierarchical structure. This hierarchy provides a communication protocol for nerve impulses from different regions.

Ripples were first observed by Cornelius Vanderwolf in 1969, after which John O’Keefe discovered place cells associated with sharp ripples [15]. Buzsaki then studied the role of sharp ripples in detail [16]. Sharp wave ripples are a compound waveform that occurs in the hippocampus. The hippocampus is an area of the brain associated with memory and learning, and sharp wave ripples are also thought to be involved in the consolidation and retrieval of memories.

Sharp wave ripples are one of the characteristic waves during slow wave sleep [17]. According to different characteristic waves, sleep can be divided into REM sleep and non-REM sleep. Non-REM sleep can be further divided into several stages, of which stages 3-4 are called slow-wave sleep (Slow-wave Sleep, SWS). Slow wave sleep has three characteristic waves, including sharp wave ripples from the hippocampus, spindle waves from the thalamus, and slow waves from the cerebral cortex.

Sharp ripples, spindles, and slow waves all occur in slow-wave sleep, and there must be some connection between them. According to Feld et al. [18], these three waves are phase-locked during sleep. Spike ripples accompany the replayed message, synchronized with the spindle excitation period of the thalamus. The spindles themselves are synchronized with the ascending phase of the cortical slow waves. During non-REM sleep, the synchronization of these three waves establishes a pathway of information from the hippocampus to the cerebral cortex.

### 4. Memory Consolidation

S Diekelmann proposed a two-stage memory consolidation model (Diekelmann & Born, 2010). During the wake stage, outside information is encoded by the cortex based on pre-existing long-term memory and the hippocampus will form temporary memory. Then during SWS sleep, the temporary memory will be reactivated in the form of replay for memory consolidation. After that, during REM sleep, synapsis will be consolidated and normalized.

As an important process of the memory system, memory consolidation is highly related to memory engrams and neural oscillations. Memory consolidation refers to the process of transforming the temporary and unstable representation of external information entering the brain into a more stable and structured long-term representation [19]. This process can be divided into two levels, namely the cellular level and the systemic level. At the cellular level, the representation of information depends on physical or chemical updates of the cell state [20–21]. and changes in the strength of synapses [19]. At the system level, the initial representation of memory will be further abstracted and transformed, especially in the offline state. This process involves communication and cooperation between several brain regions. Information flows between regions, incorporating new memories into existing ones. During this process, engram cells change. According to existing research, engrams are highly dynamic and can undergo bidirectional transformation between silent and active states. Memory consolidation is compared with the maturity of engrams. Engram cells will transform from silent into active during the consolidation process [22]. This phenomenon happens across multi-areas including the hippocampus [23–24], mPFC, and BLA [25].

As for spatial memory, Gabrielle Girardeau devised a reward-based behavioral paradigm for spatial memory consolidation [26]. Mice will be shocked either during the sharp ripple (experimental group) or after the sharp ripple (control group). The results showed that when the sharp wave ripples were disturbed, the mice performed worse on a spatial memory task. This experiment demonstrates that sharp wave ripples play an essential role in spatial memory consolidation.

### 5. Cognitive Map Theory

Regarding how the hippocampus encodes various knowledge, cognitive map theory assumes that everything could be represented by a graph structure. Everything is a relationship in the brain [16]. The psychophysical curve shows that our brains do not encode abstract value, physical value is encoded in the log form. This evidence supports the idea that the brain represents this world by relationship. In graph theory, every graph can be represented by a set of nodes and edges. In other words, we can represent any relationship in the form of a graph. Based on the abstract relationship, we can bind conceptual objects into this graph. This mechanism can be used to explain how creatures encode spatial maps and non-spatial relationships [27].

## A Unified Theory

Here we argue that sharp-wave ripples and engram cells are two different phenomena under a unified memory system from three aspects. 1) Using the cognitive map theory to unify the encoding of spatial and episodic memory. 2) Using a dynamic autoencoder conceptual model to unify static memory fingerprint and dynamic time-varying memory trace. 3) Using a spike neural network-based computation model to verify that the engram can trigger sharp-wave ripples. These three pieces of evidence resolve three divergences of sharp-wave ripples and engram cells respectively.

## Spatial and Episodic Memory Representation

One divergence between sharp-wave ripples and engram cells is that they encode different forms of memory (Figure 1). Experimental evidence shows that sequential firing and sharp-wave ripples are always coupled. And sequential firing is found in spatial memory encoding in the hippocampus area with place cells. However, engram cells are about episodic memory like a contextual-based reward or aversion memory. Lucky, sequential firing, sharp-wave ripples, and engram cells all have been observed in the hippocampus. Here I argue that

**Figure 1:**
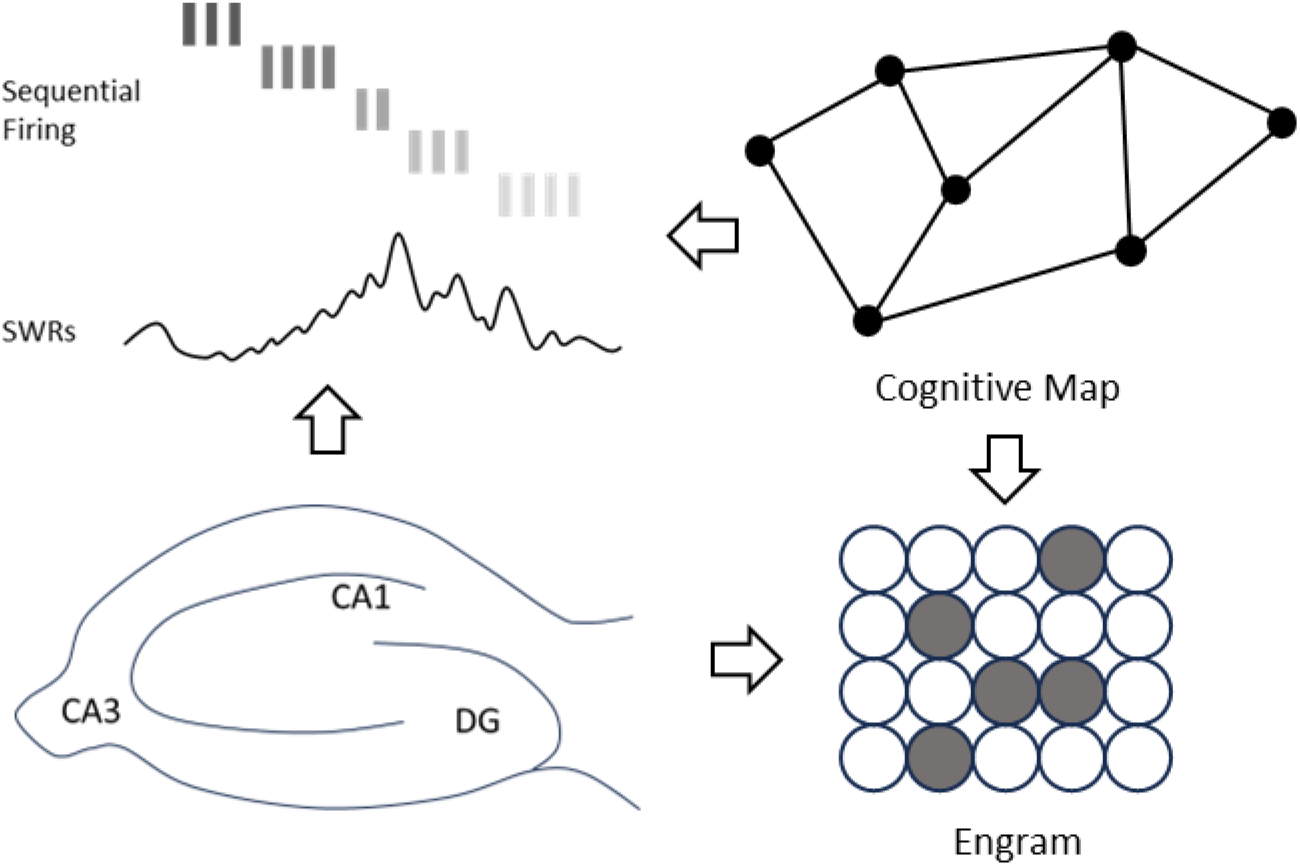
Engram and SWRs in the Hippocampus. Sharp-wave ripples are observed in the CA1 area and coupled with sequential firings. Sequential firings are coupled with spatial memory. Engram cells are observed in the DG area, which encodes episodic memory.

Animals are natively drawn on profits and avoid loss. They can learn in which environment there will have a higher chance to find food or enemies. This is called episodic memory. Animals also have spatial memory since they need to remember where is their home and where can collect resources. On the one hand, place cells are spatial in their nature, so perhaps the replay is a spatial-specific phenomenon. On the other hand, since the circuit and cellular machinery for the spatial replay are already in place, it is possible other types of memory can use this machinery if they also have a sequential pattern. This has not been sufficiently studied before.

Here we use cognitive map theory to merge that fork of spatial memory and episodic memory. We argue that spatial memory is a special case of episodic memory. As it is mentioned in the cognitive theory and Buzaski’s work [16]. The brain uses a relative relationship to encode this world. We argue that the brain uses graph structure to represent the episodic context. Spatial memory, as a spatial case, is a well-formed graph with limitations. Spatial memories are always represented as a chain (straight channel) or honeycomb-like grid (open field).

When a mouse runs on the ground, its spatial memory is a time-varying trace. This property leads to sequential firing phenomena observed in the hippocampus. However, episodic memory is much more complex. Features in context are much more enriched compared with simple coordinates in the open field. Thus, the relationship represented by the brain will be much more complex. Discover such a pattern is more computationally expensive compared with a simple chain-like structure. This might be the reason why encoding patterns about episodic memory is not discovered in the experiment yet.

According to the cognitive map theory, everything can be represented by a relation map. In every temporal frame, episodic memory can be encoded by a static graph. The graph can evolve dynamically to reflect the temporal variance of the context (As shown in Figure 2 Top Right Corner). The corresponding neuron firing pattern is more complex rather than simple sequential firing. Here I believe spatial memory is a kind of highly compressed episodic memory. In spatial memory, detailed context information is ignored and only a single coordinate is recorded. This is why we can find sequential firing in spatial memory: it is a kind of highly abstract context that only keeps coordinate information.

**Figure 2:**
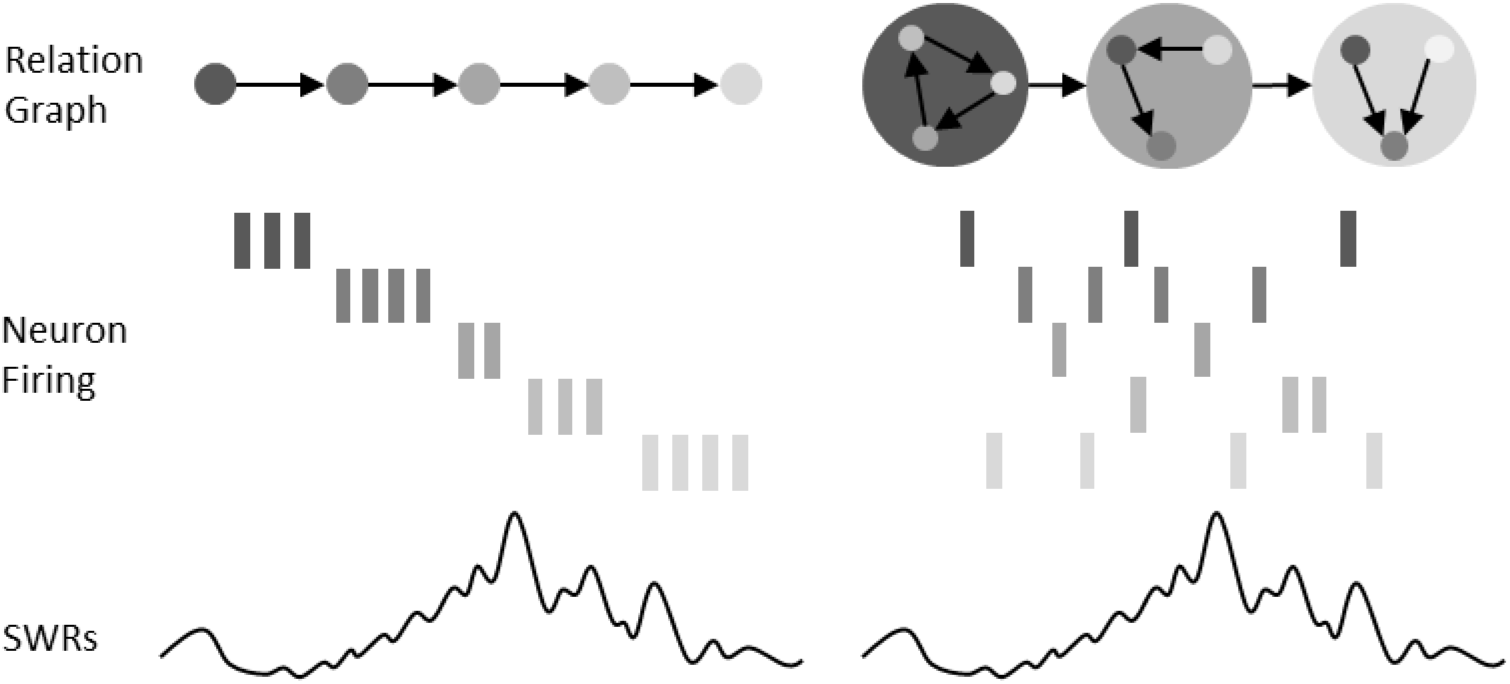
Encoding of Spatial Memory and Episodic Memory. Spatial memory is a special case of episodic memory. Every frame in the spatial memory is compressed into a single coordinate point. That’s why sequential firing can be observed.

## Dynamic Autoencoder Conceptual Model

Another divergence between sharp-wave ripples and engram cells is static fingerprint and dynamic sequence. Sharp-wave ripples are coupled with sequential firing. In the traditional experiment, engrams are a kind of spatial pattern in the DG area. The temporal property of engrams is not well-studied. In my hypothesis, the engram can trigger sharp-wave ripples. In the meanwhile, some papers believe that sharp-wave ripples can trigger sequential firing. Here I argue that it is a reasonable system to store information static in the form of spatial patterns (engram) and extract information with dynamic activities (sequential firing).

This paper believes that the working mode of forming a firing sequence through an initial stimulus is in corresponds with the phenomenon in nonlinear dynamics and the current attractor theory. Such modes enhance the reliability of neural encodings while preserving plasticity. Representing and storing sequences in the form of initial values, and using plastic manifold spaces to decompress the initial points is a faster way to represent, store, and read information efficiently. As shown in Figure 3, the encoder encodes the information and maps it to an initial point on the manifold surface. The trajectory of the initial point is described by a nonlinear dynamical system, which produces a trajectory over time. This trajectory is then sampled by the decoder, generating a sequential output. Within the framework of this model, the activation of engram cells sets the initial value. Under the description of the nonlinear dynamic equation, the initial value is decompressed into a trajectory unfolding with time, which also generates sequential information. This sequential information is the sequential firing observed in the experiment.

**Figure 3.**
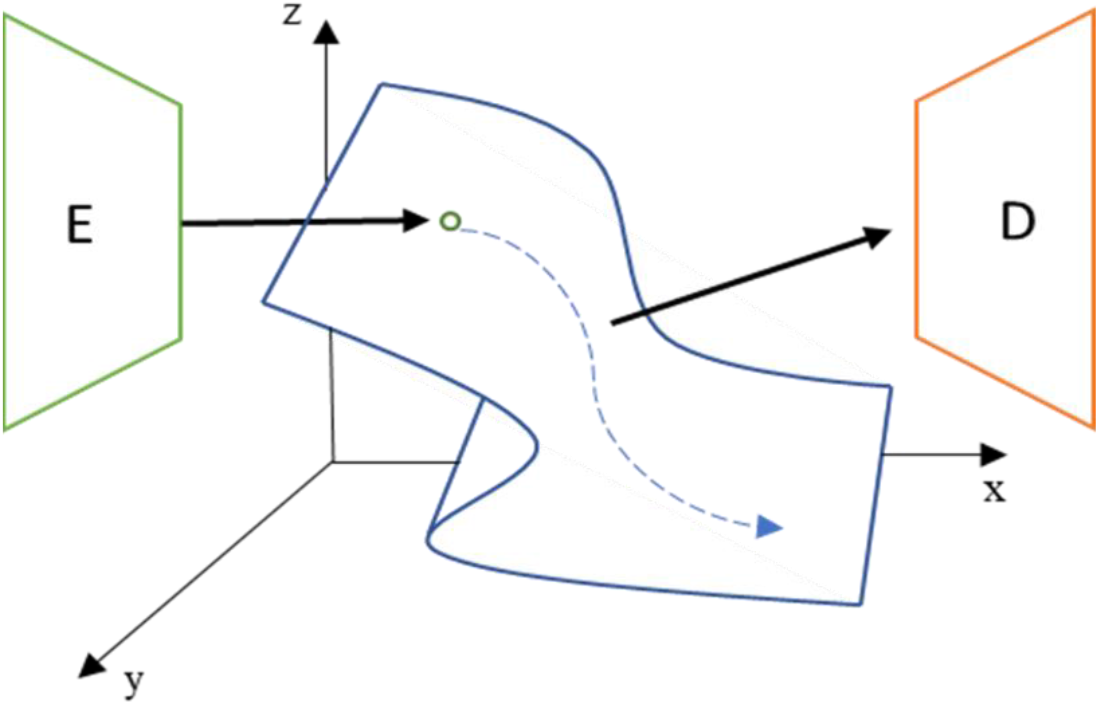
A Dynamic Autoencoder Model of The Hippocampus. In previous work, it was proposed that the hippocampus can be considered an autoencoder, and the theory is extended here in conjunction with a dynamic autoencoding model. The encoder sets an initial value on the manifold, and the subsequent movement of the initial coordinates is described by a nonlinear dynamical system. Its time-varying motion trajectory is sampled by the decoder to generate a sequential output.

This conceptual model is supported by many studies. Benna et al. [28] argue that the entorhinal cortex and hippocampus constitute an autoencoder. Jaffe et al. [29] proposed a task-dependent dynamic self-variational encoder [30] to model human behavior. The encoding from an initial point to a trajectory is a universal computational principle in the brain. In the field of motion control, Wang et al. [31] mentioned that the motor cortex generates temporal motion commands according to the dynamic evolution of the trajectory by setting the initial value. In the field of value coding and decision-making, Okazawa et al. [32] found that population neuron coding forms a manifold space in the lateral parietal cortex. In the computational model of working memory, Bi et al. [33] believed that the track length encoded the delay time between different stimulus signals. This paper argues that memory consolidation and representation can also be included under the framework of this universal computational principle.

## Spike Neural Network-based Computational Model

Biological experiments are constrained by equipment, instruments, reagents, and experimental methods. The verification and observation of new phenomena require a long experimental cycle and may involve many interference factors. Therefore, starting from the existing experimental results, we leveraged a computational model to verify the hypothesis that engram cells can trigger sharp-wave ripples.

### 1. Tri-synaptic Pathway in the Hippocampus

There are a large number of excitatory and inhibitory cells in the CA1 and CA3 regions, and the calculation model also shows that the interaction of excitatory and inhibitory cells is required to construct the attractor model. Excitatory and inhibitory cells play a role in homeostasis. As shown in Figure 4, there are a large number of projections from the DG area to CA3 and then to CA1, forming a tri-synaptic loop. The sharp wave-ripples phenomenon is similar to the damped oscillation phenomenon. Thus, activation of engrams in the DG region may introduce an additional input in the CA3 region [34]. Inputs to the DG disrupt the balance between excitatory and inhibitory neurons in the CA, producing ripples. And the forward sharp wave will also induce the chain firing of neurons along the way, realizing the phenomenon of sequential firing. In short, the sharp-wave ripples might be an energy dissipation phenomenon in the excitatory-inhibition balanced system.

**Figure 4:**
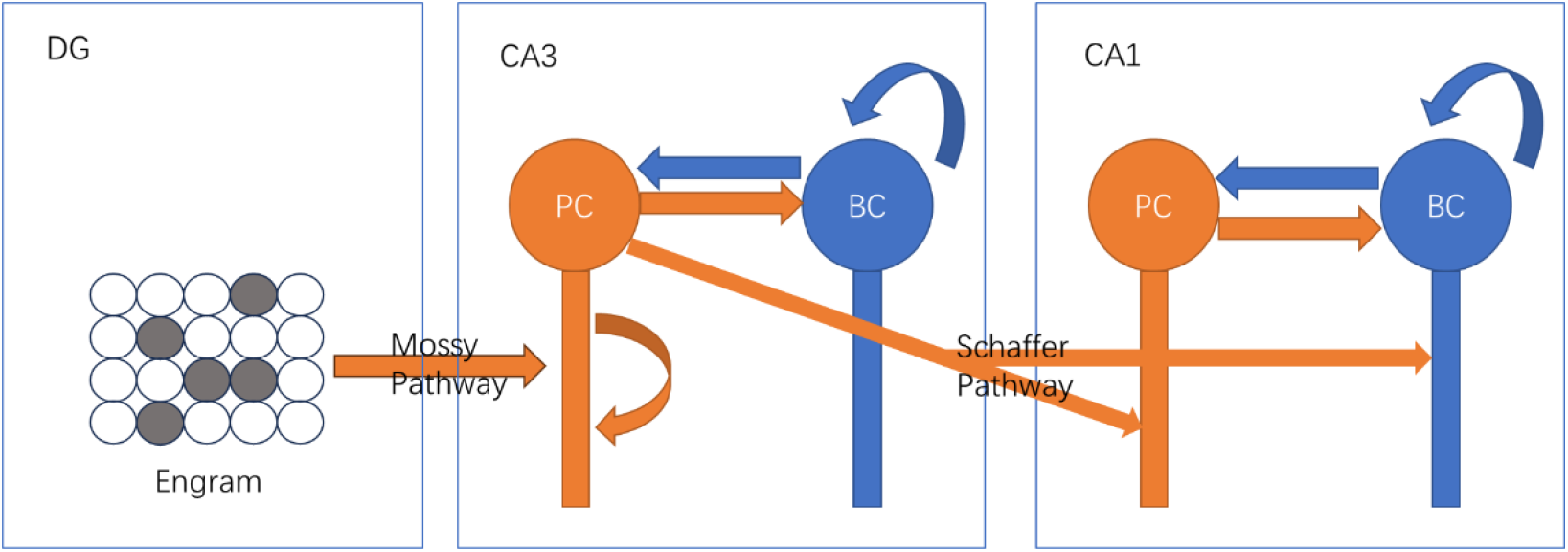
Tri-synaptic Pathway. The DG-CA3-CA1 connectivity pathway is referred to as the tri-synaptic pathway. A multi-compartment model is used for the CA3 and CA1 regions. Arrows represent projection relationships. Orange represents inhibitory and blue represents excitatory. In the multi-compartment model, the circle represents the soma and the rectangle represents the axon.

### 2. Energy View of the E-I Balanced System

As shown in Equation 1, the equation is based on the Hodgkin-Huxley model. This model describes the relationship between the neuronal membrane potential and the input current. The CA3 and CA1 regions are considered as an entirety. From the perspective of charge, the only input quantity of the whole system is the drive current *I_drive_* from the DG region reaches the CA3 region through the Mossy pathway. The charge brought by the drive current will be stored by the neuron in the form of membrane potential and released in the form of the action potential. From this perspective, the whole model of the hippocampus can be thought of as a capacitor.

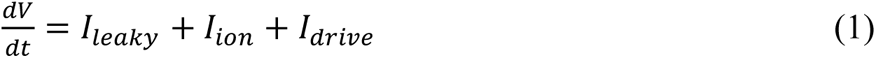

Both CA3 and CA1 contain two groups of neurons, excitatory neurons and inhibitory neurons. Simply thinking about the relationship between the two groups of neurons, we can know that they constitute a self-feedback stable system. The self-feedback system can be simply extended to the analysis of charge storage. Taking the CA1 region as an example, the input of the Schaffer pathway leads to an increase in the membrane potential of excitatory neurons. An increase in the membrane potential of excitatory neurons leads to an increase in the membrane potential of inhibitory neurons, thereby inhibiting the increase in the membrane potential of excitatory neurons. The increased energy can be released in the form of higher firing frequency of both excitatory and inhibitory neurons. In the meantime, the whole system maintains a stable state. In other words, the two groups of neurons work in antagonism. But something will change in the system. A higher rate of action potential firing leads to more charges being consumed. But the entire system maintains a dynamic balance, and the amount of charges will not accumulate infinitely.

From the perspective of energy, sharp-wave ripples are the result of the energy explosion. Therefore, a reasonable deduction is that the generation of sharp-wave ripples must be accompanied by a decrease in the amount of stored charge in the neuron population. It has also been found that the synchronous firing of CA1 inhibitory neurons can induce the generation of sharp-wave ripples [35]. This phenomenon can also be explained from the perspective of charge storage. If the CA1 area is regarded as a capacitor, it is assumed that its input from CA3 does not change but an external input enhances the firing of inhibitory neurons in the CA1 region. This leads to a decrease in the overall action potential firing rate in the CA1 region, resulting in an increase in stored charge. Sudden cessation of external input in the CA1 region will cause excitatory neurons to fire in bursts. By releasing excessive charge, sharp-wave ripples will be generated. Based on this mechanism, sharp-wave ripples can also be induced by a sudden increase in external input, and this phenomenon has also been successfully observed in a computational model.

### 3. Modeling Results

#### Sharp-Wave Ripples

This paper examined sharp-wave ripples on a computational model of the hippocampus. Please refer to supplementary materials for more details about the computational model. By simulating neuronal activity in the hippocampus over 4 s, we recorded the membrane potential of neurons in the CA1 region. Local field potentials were calculated based on membrane potentials. Energy density measurements are performed on the field potential to identify sharp ripple events. The result is shown in Figure 5.

**Figure 5:**
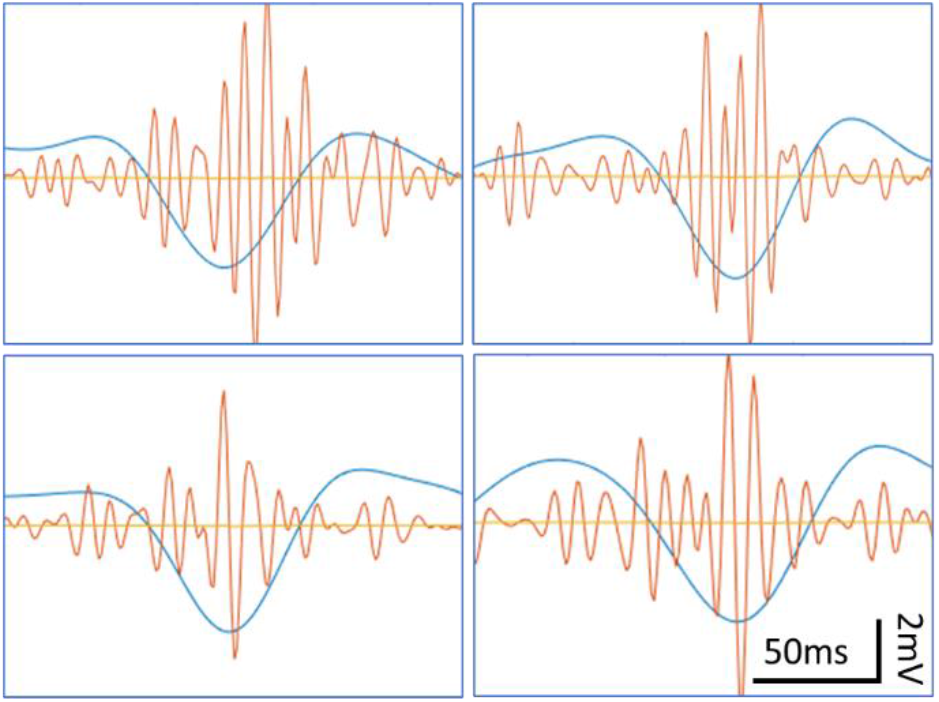
Sharp-Wave Ripples in the Computational Model. Sharp-wave ripples detected in computational models of the hippocampus. The blue curve is a sharp wave, which is obtained by bandpass the local field potential signal from 5 Hz to 20 Hz. The orange curve is the ripple, which is obtained by bandpass the field potential signal from 80 Hz to 180 Hz. The yellow horizontal line is the baseline, representing zero potential.

#### Trigger Sharp-Wave Ripples by Temporary Enhanced Input

The paradigms above provide a possible mechanism for the generation of sharp-wave ripples from the perspective of neuron population charge. Based on this mechanism, a reasonable inference can be drawn: the disturbance brought by the input of the DG area disrupts the balance between the excitatory and inhibitory neurons in the CA1 and CA3 areas, resulting in sharp-wave ripples. To test this hypothesis, the following experiments were performed in a computational model of the hippocampus. As shown in Figure 4, DG-CA3-CA1 constitutes a tri-synaptic pathway. Activation of engram cells in the DG will lead to temporarily enhanced input of the Mossy pathway. This paper simulates the activation of engram cells in the DG region by artificially changing the input current intensity of the CA3 region. Through experiments, it was found that the temporary enhancement of the Mossy pathway can induce the generation of sharp-wave ripples. As shown in Figure 6, the drive current in the CA3 area is shown by the orange curve, and it will switch between 2.3*uA*∼3*uA* in a period of 1 second. The blue curve is the local field potential in the CA1 area.

**Figure 6:**
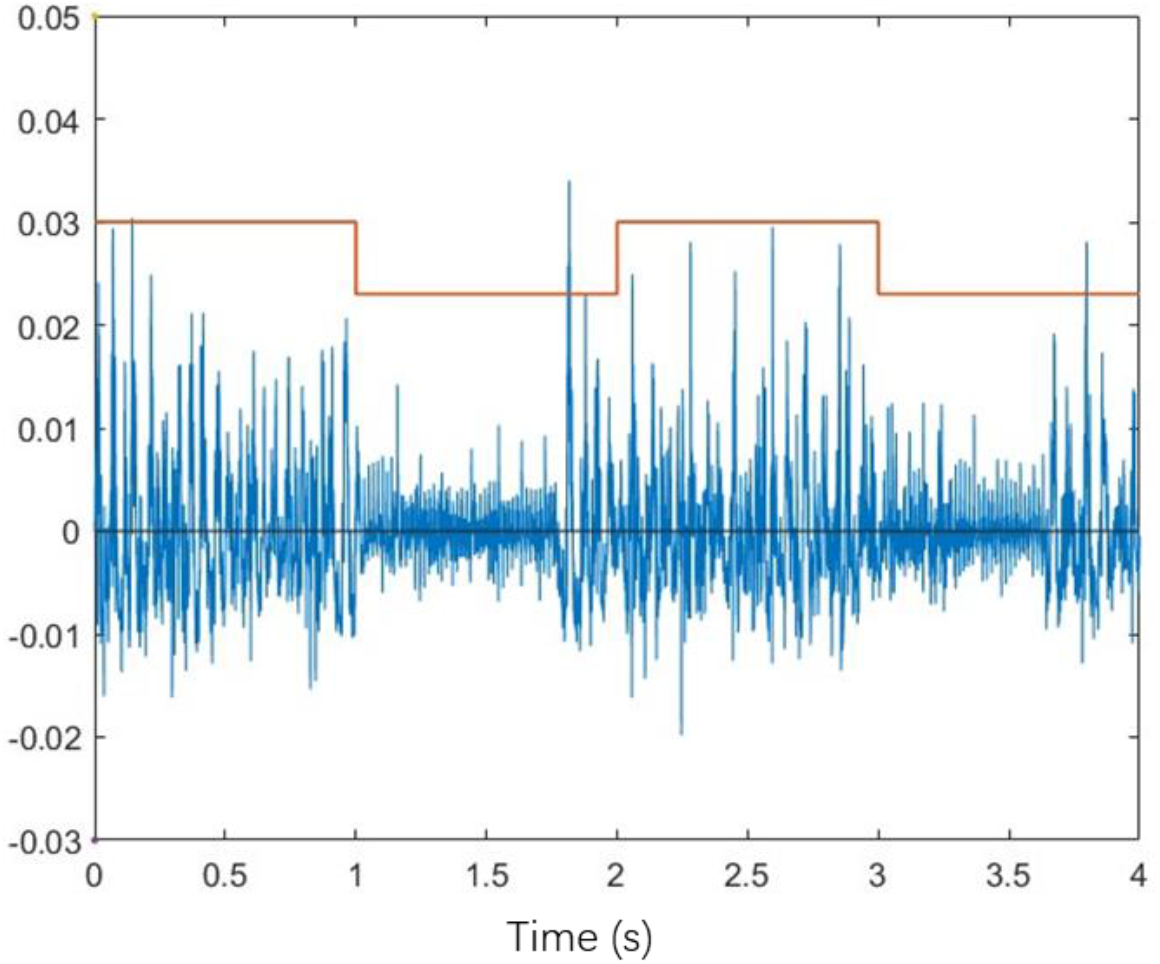
Local Field Potential in CA1 Driven by DG Input. The blue line represents the local field potential of the CA1 region, and the orange line represents the input drive current of the CA3 region. The drive current fluctuates between 2.3∼3 *μA* with a period of 1s to simulate the temporary enhancement of DG input.

As the input from the DG region switches between augmented and normal input on a one-second cycle, the probability of sharp-ripple events in the CA1 region changes. In this paper, the activity of the hippocampal region was simulated for 4 s each time, and after 40 repetitions, the frequency of sharp-wave ripples in the CA1 region was detected. The results are shown in Figure 7, the occurrence frequency of sharp ripples in the enhanced input period is significantly higher than that in the normal input period. This result supports the hypothesis that temporarily enhanced input to the DG region can induce sharp-wave ripples. Another possibility is that the enhanced input itself causes more sharp-wave ripples. After experimenting with the computational model, it is found that the absolute value of the drive current does not determine the frequency of sharp ripples. The key factor is the step switch of the drive current.

**Figure 7:**
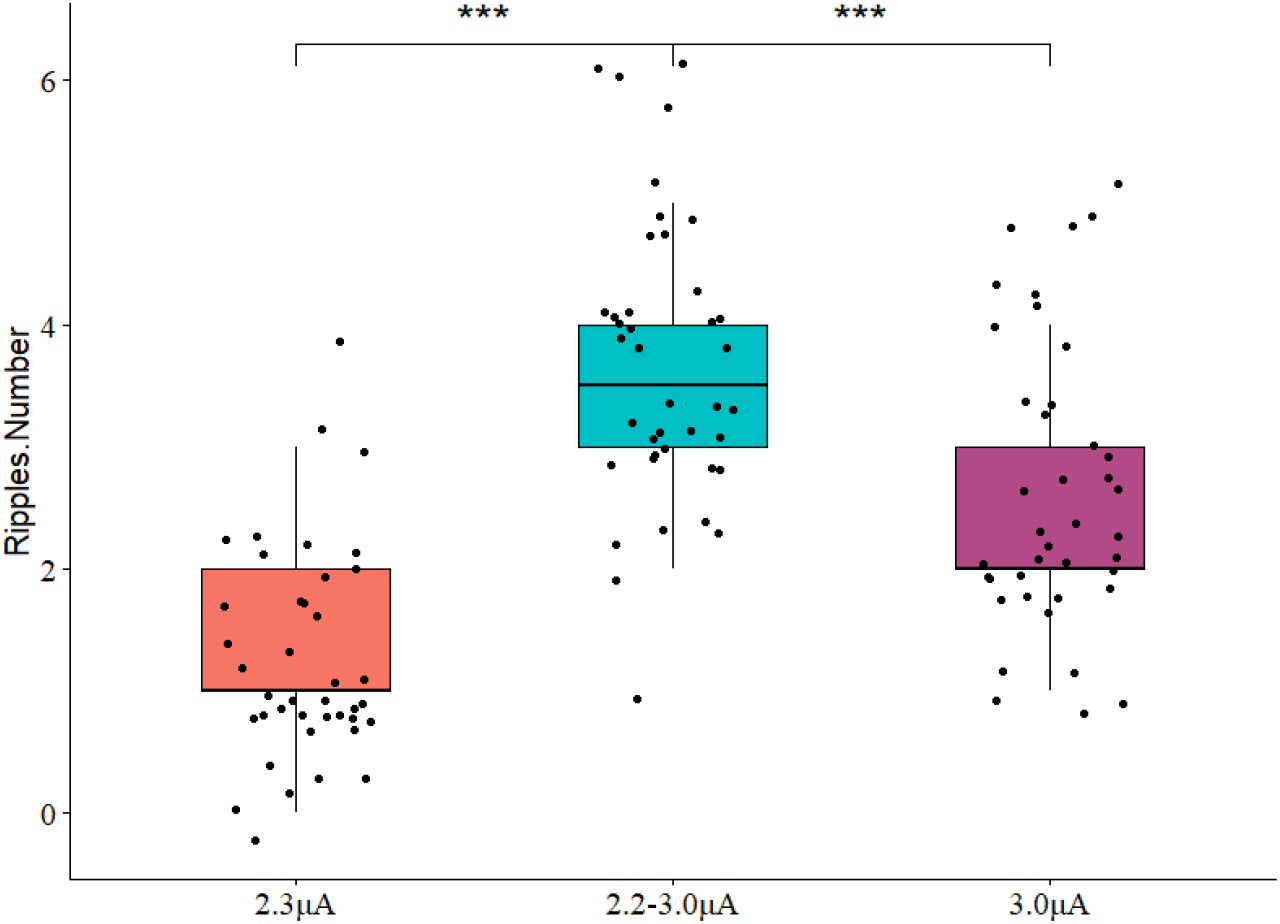
Temporary Enhanced DG Input Triggers Sharp-Wave Ripples. Stats graph and hypothesis testing proved that temporary enhancement of DG input can trigger sharp-wave ripples in the CA1 area. Run 4 s simulation 40 times. ***p < 0.001, Wilcox test.

## Conclusion & Future Work

The hippocampus is proven to play a key role in memory encoding, consolidation, and retrieval. Place cells fire in sequence to encode spatial memory. Previous studies show that ripples, spindles, and slow waves will synchronously transform information between the hippocampus and cortex.

The engram cell theory was first proposed by Semon, and Tonegawa further developed this theory, which was supported by experimental evidence at three levels: observation, loss of function, and gain of function. The long-term enhancement theory has revealed the neural mechanism of memory and learning at the cellular level. Engram formation is dependent on long-term potentiation, an increase in dendritic spines, and an increase in cellular excitability. Multiple brain regions are involved in this process, including the hippocampus, medial prefrontal cortex, and basolateral amygdala. Various regions complete this process through cooperation and produce a joint effect. However, there are still many questions to be answered. Questions for further research include how to achieve retrieval, consolidation, and representation of different memories in a unified system. At the same time, we also need to explore the information exchange mode of engram cells in different brain regions and understand the role of engram cells in different types of memory. To better study these topics, we need more advanced neural recording technology and more refined neural manipulation methods. Future studies will use higher temporal and spatial resolution methods to explore the changes of engram cells during learning and the formation of different types of memories.

Like wave-particle duality, static engram cells and dynamic sequential firing should be regarded as two representations of the same memory system. If the engram cell theory is regarded as a static memory mark, the sharp-wave ripples and the accompanying sequential replay can be regarded as a dynamic representation with highly temporal characteristics. The main research on the replay phenomenon has focused on spatial memory, while the research on engram cells has focused on episodic memory. But this paper thinks that these two forms should be equivalent and have a causal relationship.

This paper analyzed the generation mechanism of sharp-wave ripples from the perspective of computational neuroscience. Combining the charge perspective and the self-feedback mechanism, the possible reasons for the sharp-wave ripples are expounded. Based on this mechanism, this paper proposes the hypothesis that the activation of engram cells can induce sharp wave ripples, and has been verified in a computational model. In addition, this hypothesis can be verified by observing the engram and sequence firing at the same time. The verification experiment needs to be conducted by other researchers. At the model level, I hope to answer the following questions: 1) Different engram cells correspond to different memories, how to make different engram cells activate different sequence replays in the model? 2) How to add new memory engrams to the existing network model? 3) How do old engrams affect the representation of new engrams in the hippocampus?

## Supplementary Materials

The temporal dynamics of neurons are described by the Hodgkin–Huxley model. Specifically, the two-compartment model from the paper by Ramirez-Villegas et al. [36] was used, consisting of two parts, the soma, and the axon. The equation for excitatory pyramidal cells [37] is shown in Equation 1 and for inhibitory interneurons [36] is shown in Equation 2. where *C*_*m*_ represents the membrane capacitance, V represents the membrane potential, I represents the transmembrane current, and *g*_*C*_ represents the conductance between compartments. Subscript s stands for cell body, d stands for axon, *L*_*s*_ stands for leakage current, *syn* stands for synaptic current, p=0.5. The calculation of the field potential also follows the approach of Ramirez-Villegas et al. [36].

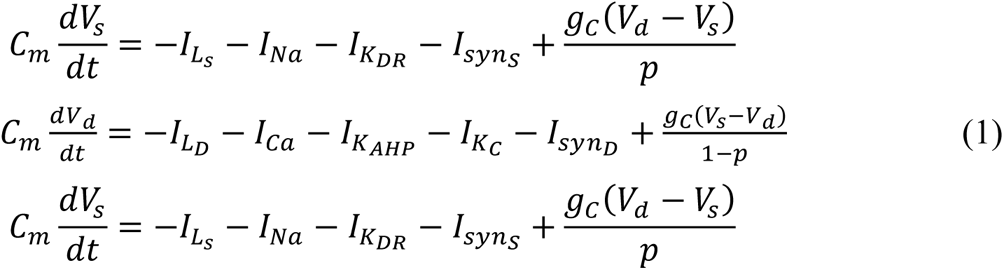

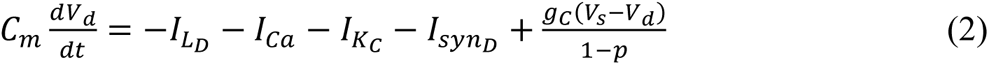

The synaptic model is shown in Equation 3. Among them, *G*_*th*_ represents the synaptic threshold, and only when the membrane potential is greater than this threshold can it produce an effect at the synapse. *τ*_*syn*_ represents the decay time constant. *φ* characterizes synaptic strength, determined by pre- and post-synaptic neuron types. *E*_*syn*_ stands for reversal potential. The specific values are shown in the attached Table 1 and attached Table 2.

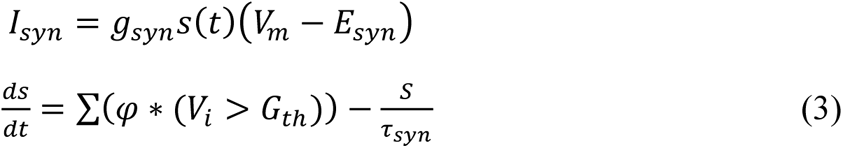

**Table 1:** Synaptic Strength of Hippocampus Model.

| Synapse Type | Synaptic Strength |
| --- | --- |
| E-E (CA3) | 2.5 |
| E-I (CA3) | 1.25 |
| I-E (CA3) | 80. |
| I-I (CA3) | 20. |
| E-I (CA1) | 0.8 |
| I-E (CA1) | 60. |
| I-I (CA1) | 30. |
| E-E (Schaffer) | 2. |
| E-I (Schaffer) | 0.8 |

Not listed in the table means that there is no projective relationship.

**Table 2:**
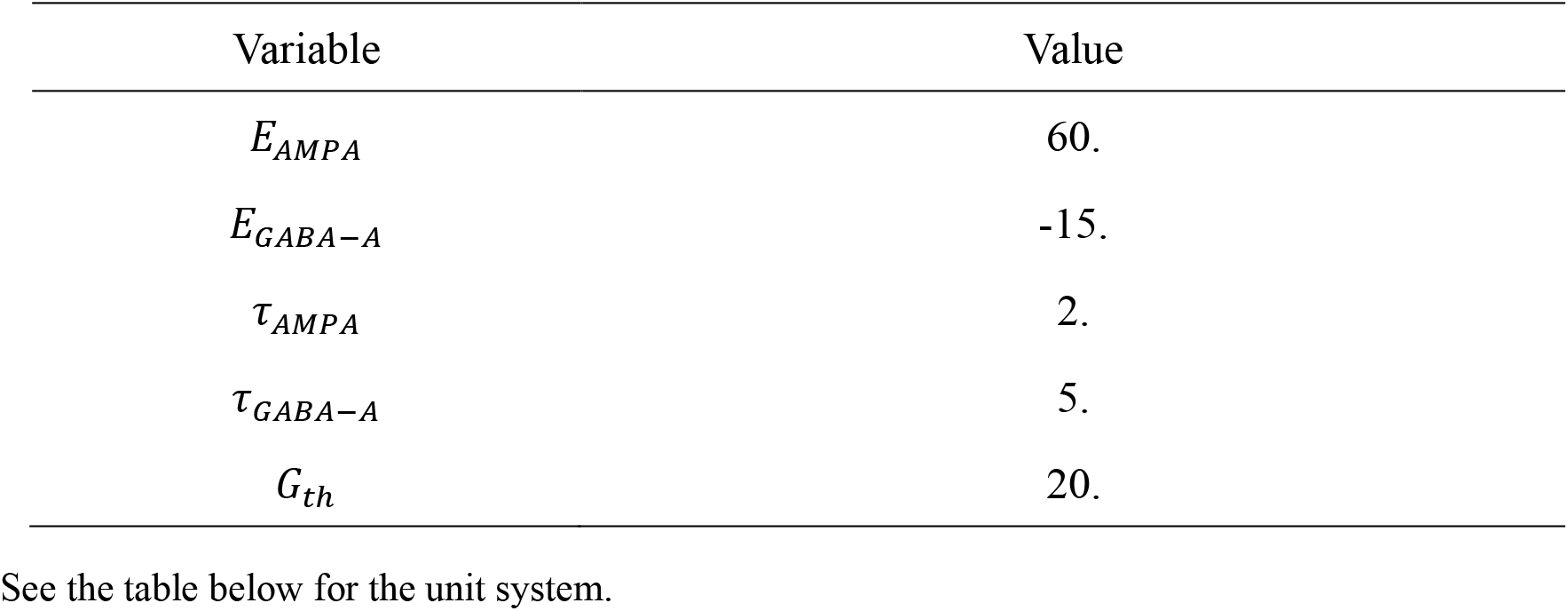
Synaptic Parameters of Hippocampus Model.

The numerical simulation method adopts the fourth-order Runge-Kutta method, and its form under the autonomous system is shown in Equation 4. Where *f*(∗) is the objective function of the numerical simulation, x is the output result, and *δ*_*t*_ is the time step. The error amount of the numerical simulation is determined by *δ*_*t*_, which is 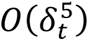. The *δ*_*t*_ = 0.02 *ms* is used in this paper.

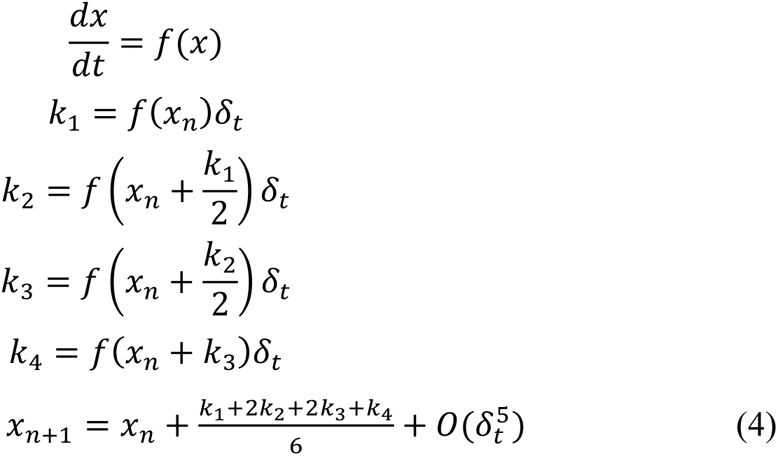

Connections within neuronal populations and between CA1 and CA3 are described probabilistically. It can be divided into two types: one is the connection probability that is inversely proportional to the distance (as shown in Equation 1-5), and the other is the uniform connection probability that is independent of distance. The projections of CA3 inhibitory neurons are distance-independent uniform projections, and the others are distance-dependent connection probabilities. See the attached Table 3 for specific parameters. See attached table 4 for the relationship between model scale and spatial location, n represents the number of neurons, and d represents the distance between neurons.

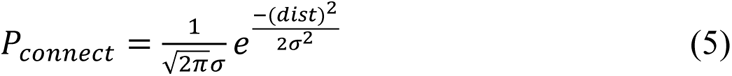

**Table 3:** Projection Probability of Hippocampus Model.

| Variable |  |
| --- | --- |
| $\sigma_E$ | 1. |
| $\sigma_I$ | 0.1 |
| $\sigma_{CA3-CA1}$ | 1.2 |

$\sigma_E$ and $\sigma_g$ apply to both CA1 and CA3.

**Table 4:** Scale and Spatial Distribution of Hippocampus Model.

| Variable | Value |
| --- | --- |
| $n_E$ | 135 |
| $n_I$ | 15 |
| d | 0.01 mm |
| $d_{CA3-CA1}$ | 1.0 mm |

n represents the number of neurons, and d represents the distance between neurons.

The unit system is shown in Attached Table 5. If there is no special description, this unit system is used by default. For variables not in this table, it is dimensionless.

**Table 5:**
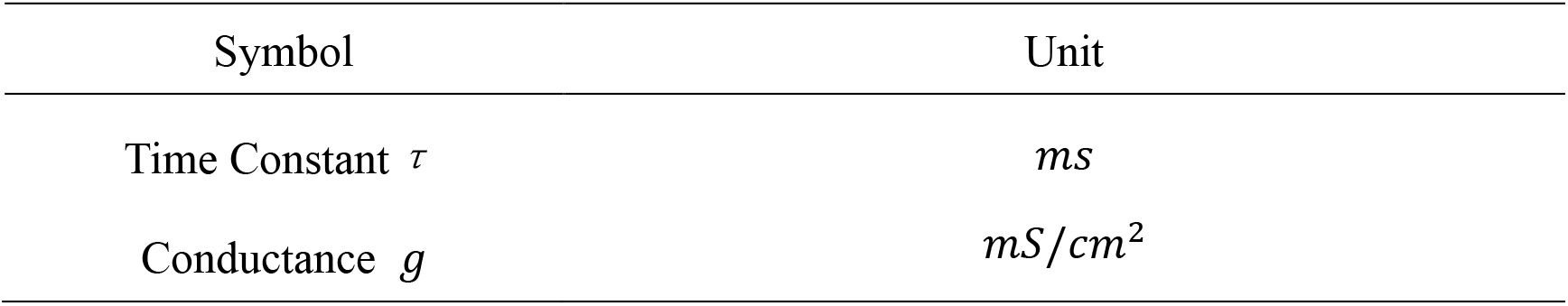
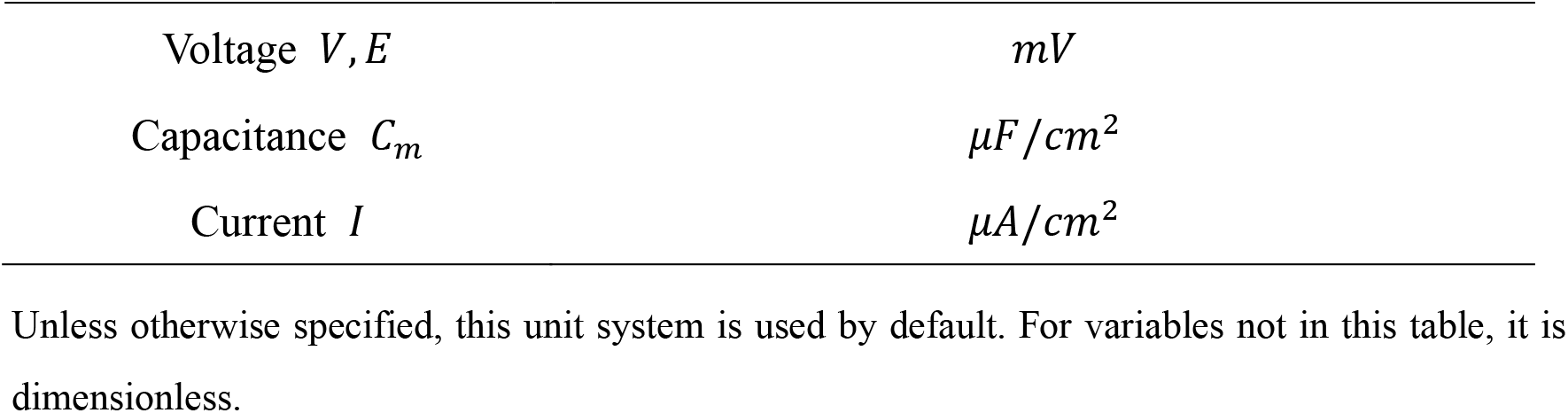
Unit System of Computational Model.

| Symbol | Unit |
| --- | --- |
| Time Constant $\tau$ | ms |
| Conductance $g$ | mS/cm<sup>2</sup> |
| Voltage $V, E$ | $mV$ |
| Capacitance $C_m$ | $\mu F/cm^2$ |
| Current $I$ | $\mu A/cm^2$ |

Unless otherwise specified, this unit system is used by default. For variables not in this table, it is dimensionless.

## Reference

[1] Tonegawa S, Liu X, Ramirez S, et al. Memory engram cells have come of age[J]. Neuron, 2015, 87(5): 918–931.

[2] Josselyn S A, Tonegawa S. Memory engrams: Recalling the past and imagining the future[J]. Science, 367.

[3] Kim, J. J, Fanselow, et al. Modality-specific retrograde amnesia of fear[J]. Science, 1992.

[4] Furber S B, Bainbridge W J, Cumpstey J M, et al. Sparse distributed memory using N-of-M codes[J]. Neural Networks the Official Journal of the International Neural Network Society, 2004, 17(10):1437–1451.

[5] Choi J H, Sim S E, Kim J I, et al. Interregional synaptic maps among engram cells underlie memory formation[J]. Science, 2018, 360(6387):430.

[6] Tanaka K, Pevzner A, Hamidi A, et al. Cortical representations are reinstated by the hippocampus during memory retrieval[J]. Neuron, 2014, 84(2):347–354.

[7] Goshen I, Brodsky M, Prakash R, et al. Dynamics of Retrieval Strategies for Remote Memories[J]. Cell, 2011, 147(5):1197–1197.

[8] Kheirbek M A, Drew L J, Burghardt N S, et al. Differential Control of Learning and Anxiety along the Dorsoventral Axis of the Dentate Gyrus[J]. Neuron, 2013, 77(5):955–968.

[9] Ryan T J, Roy D S, Pignatelli M, et al. Memory. Engram cells retain memory under retrograde amnesia[J]. Science, 2015, 348(6238):1007-1013.

[10] Liu X, Ramirez S, Pang P T, et al. Optogenetic stimulation of a hippocampal engram activates fear memory recall[J]. Nature, 2012, 484(7394): 381-385.

[11] Ramirez S, Liu X, Macdonald C J, et al. Activating positive memory engrams suppresses depression-like behaviour[J]. Nature, 2015, 522(7556):335–339.

[12] Ramirez S, Liu X, Lin P A, et al. Creating a false memory in the hippocampus[J]. Science, 2013, 341(6144): 387–391.

[13] Vetere G, Tran L M, Moberg S, et al. Memory formation in the absence of experience[J]. Nature Neuroscience, 2019, 22(6): 933–940.

[14] Ji D, Wilson M A. Coordinated memory replay in the visual cortex and hippocampus during sleep[J]. Nature Neuroscience, 2007, 10(1):100–107.

[15] Maier N, Draguhn A, Schmitz D, et al. Fast network oscillations in the hippocampus[J]. e-Neuroforum, 2013, 19(1): 1–10.

[16] György Buzsáki M D. The brain from inside out[M]. Oxford University Press, 2019.

[17] Chambers A M, Payne J D. The Memory Function of Sleep[M]. John Wiley & Sons, Ltd, 2015.

[18] Feld G B, Born J. Sculpting memory during sleep: concurrent consolidation and forgetting[J]. Current Opinion in Neurobiology, 2017, 44:20–27.

[19] Lechner H A, Squire L R, Byrne J H. 100 Years of Consolidation— Remembering Müller and Pilzecker[J]. Learning & Memory, 1999, 6(2):77.

[20] Lesburgueres E, Gobbo O L, Alaux-Cantin S, et al. Early Tagging of Cortical Networks Is Required for the Formation of Enduring Associative Memory[J]. Science, 2011, 331(6019):924-928.

[21] Miller C A, Gavin C F, White J A, et al. Cortical DNA methylation maintains remote memory[J]. Nature Neuroscience, 2010, 13(6): 664–666.

[22] Susumu, Tonegawa, Mark D, et al. The role of engram cells in the systems consolidation of memory.[J]. Nature Reviews. Neuroscience, 2018.

[23] Kitamura T, Ogawa S K, Roy D S, et al. Engrams and circuits crucial for systems consolidation of a memory[J]. Science, 2017, 356(6333):73–78.

[24] Maren S, Aharonov G, Fanselow M S. Neurotoxic lesions of the dorsal hippocampus and Pavlovian fear conditioning in rats[J]. Behavioural Brain Research, 1997, 88(2):261–74.

[25] Kitamura T, Ogawa S K, Roy D S, et al. Engrams and circuits crucial for systems consolidation of a memory[J]. Science, 2017, 356(6333):73–78.

[26] Girardeau G, Benchenane K, Wiener S I, et al. Selective suppression of hippocampal ripples impairs spatial memory[J]. Nature Neuroscience, 2009, 12(10):1222–1223.

[27] Whittington, J. C. R., Muller, T. H., Mark, S., Chen, G., Barry, C., Burgess, N., & Behrens, T. E. J. (2020). The Tolman-Eichenbaum Machine: Unifying Space and Relational Memory through Generalization in the Hippocampal Formation. Cell, 183(5), 1249–1263.e23. https://doi.org/10.1016/j.cell.2020.10.024

[28] Benna M K, Fusi S. Place cells may simply be memory cells: Memory compression leads to spatial tuning and history dependence[J]. Proceedings of the National Academy of Sciences, 2021, 118(51): e2018422118.

[29] Jaffe P I, Poldrack R A, Schafer R J, et al. Modelling human behaviour in cognitive tasks with latent dynamical systems[J]. Nature Human Behaviour, 2023: 1–15.

[30] Girin L, Leglaive S, Bie X, et al. Dynamical variational autoencoders: A comprehensive review[J]. arXiv preprint arXiv:2008.12595, 2020.

[31] Wang T, Chen Y, Cui H. From parametric representation to dynamical system: shifting views of the motor cortex in motor control[J]. Neuroscience Bulletin, 2022, 38(7): 796–808.

[32] Okazawa G, Hatch C E, Mancoo A, et al. Representational geometry of perceptual decisions in the monkey parietal cortex[J]. Cell, 2021, 184(14): 3748–3761. e18.

[33] Bi Z, Zhou C. Understanding the computation of time using neural network models[J]. Proceedings of the National Academy of Sciences, 2020, 117(19): 10530–10540.

[34] Guan J S, Jiang J, Hong X, et al. How Does the Sparse Memory “Engram” Neurons Encode the Memory of a Spatial–Temporal Event?[J]. Frontiers in Neural Circuits, 2016, 10.

[35] Stark E, Roux L, Eichler R, et al. Pyramidal cell-interneuron interactions underlie hippocampal ripple oscillations[J]. Neuron, 2014, 83(2): 467–480.

[36] Ramirez-Villegas J F, Willeke K F, Logothetis N K, et al. Dissecting the synapse and frequency-dependent network mechanisms of in vivo hippocampal sharp wave-ripples[J]. Neuron, 2018, 100(5): 1224–1240. e13.

[37] Pinsky P F, Rinzel J. Intrinsic and network rhythmogenesis in a reduced Traub model for CA3 neurons[J]. Journal of Computational Neuroscience, 1994, 1: 39–60.

